# *In Vitro* Probiotic Potential of Hemophilin-producing Strains of *Haemophilus haemolyticus*

**DOI:** 10.1101/2020.01.02.893487

**Authors:** Brianna Atto, Roger Latham, Dale Kunde, David Gell, Stephen Tristram

**Affiliations:** School of Health Sciences, University of Tasmania, Launceston, TAS, Australia; School of Medicine, University of Tasmania, Hobart, TAS, Australia

## Abstract

Non-typeable *Haemophilus influenzae* (NTHi) is a leading causative organism of opportunistic respiratory tract infections, including otitis media and acute exacerbations of chronic obstructive pulmonary disease. Despite the enormous disease burden associated with NTHi infections, there are currently no effective prevention strategies, and the rapid development of antibiotic resistance is compromising treatment.

We previously discovered *Haemophilus haemolyticus* (Hh) strains capable of producing haemophilin (HPL), a heme-binding protein that restricts NTHi growth by limiting its access to an essential growth factor, heme. Thus, these strains may have utility as a probiotic therapy against NTHi infection by limiting colonization, migration and subsequent infection in susceptible individuals. Here, we have assessed the feasibility of this approach by *in vitro* competition assays between NTHi and Hh strains with varying capacity to produce HPL. HPL-producing strains of Hh exhibited enhanced growth and consistently outcompeted NTHi compared to Hh strains unable to produce the protein. This competitive advantage was maintained over a period of six days, culminating in the complete eradication of NTHi. Expression analysis of *HPL* during competition coincided with the NTHi-inhibitory capacity of HPL-producers, confirming that inhibition was mediated by the presence of HPL.

Together, results suggest that natural levels of HPL production by Hh are sufficient to limit NTHi’s access to heme, even under excess heme conditions unlikely to be encountered *in vivo.* Further investigation is required to determine the protective capacity of HPL-producers *in vivo* and their ability to interrupt NTHi colonization of host cells.

## INTRODUCTION

The bacterium non-typeable *Haemophilus influenzae* (NTHi) is commonly associated with upper respiratory tract (URT) colonization in healthy adults (1). However, migration to other sites in the respiratory tract frequently occur in children, the elderly and individuals with underlying respiratory diseases; making NTHi a leading cause of mucosal infections (2–4). In particular, enormous global morbidity is attributed to otitis media (OM) and exacerbations of chronic obstructive pulmonary disease (COPD), which are accompanied by long-term health complications and considerable mortality, respectively (5, 6). NTHi has also gained attention as an increasingly important cause of invasive infections (7–9).

There are currently no effective vaccination strategies for the prevention of NTHi infections and treatment has been complicated by the rapid development of antibiotic resistance to first- and second-line antibiotics. Resistance is predominantly mediated by β-lactamase production (10); however, the emergence and spread of β-lactamase-negative, ampicillin-resistant strains in many regions of the world is of substantial concern with treatment failure also being reported in response to macrolides (11–14) and fluoroquinolones (15–17).

NTHi infection is preceded by successful colonization of the URT and survival in this environment relies on the bacterium’s ability to acquire the vital growth factor, heme (18). There is also evidence to suggest heme-acquisition genes are important modulators of NTHi virulence factors (19), demonstrated by the increased prevalence in disease-causing strains from the middle ear, compared to colonizing throat strains (18). Deletion of multiple genes related to haem-iron scavenging, utilization and regulation has been shown to significantly reduce NTHi virulence, disease severity and duration in animal models of OM (20, 21). Similarly, an isogenic mutant of two haem-acquisition pathways was unable to sustain bacteraemia or produce meningitis in a rat model of invasive disease (22). Thus, haem-acquisition pathways represent potentially high value targets for the development of novel therapies for the eradication NTHi from the respiratory tract (23, 24).

NTHi is particularly susceptible to heme restriction as it is incapable of heme synthesis and relies solely on scavenging heme from the host, either in the form of free-heme or bound to host carrier molecules (20, 25–27). Evidence from our laboratory suggests that closely related commensals may present a competitive challenge for heme acquisition in the URT. Previously, we discovered *Haemophilus haemolyticus* (Hh) strains that exhibited inhibitory activity against NTHi (28, 29). Further investigation revealed this inhibition was mediated by the production of a heme-binding protein, haemophilin (HPL) that restricted NTHi growth by limiting its access to heme (29). Thus, these strains may have utility as a probiotic therapy against NTHi infection by limiting colonization, migration and subsequent infection in susceptible individuals. Here, we aim to determine the feasibility of the probiotic approach by assessing *in vitro* competition between NTHi and HPL-producing strains of Hh.

## METHODS

### Bacterial growth conditions

#### Bacterial Strains

Hh strains used in this study have previously been isolated and screened for the *HPL* ORF (28, 29) An *HPL* knockout (BW1^*HPL*−^) of a model HPL-producing strain of Hh (Hh-BW1) was constructed using insertional inactivation as previously described (29).

NTHi and Hh isolates were propagated from liquid nitrogen frozen glycerol stock, followed by two overnight passages on chocolate agar (CA) at 37°C with 5–10% CO_2_ prior to experimentation. Strains were grown in supplemented Tryptone Soy Broth (sTSB), which consisted of tryptone soy broth (TSB) (Oxoid Ltd., Basingstoke, UK) supplemented with 2% (v/v) Vitox® (Oxoid Ltd) and 15 μg mL^−1^ of porcine haematin (ferriprotoporphyrin IX hydroxide, Sigma-Aldrich). Exposure to non-growth conditions was minimized by maintaining suspensions and diluents at 37°C in heat block with sand or benchtop incubator.

#### Propagation of heme-replete populations for growth experiments

Strains were also propagated under heme-replete conditions prior to competition to replenish bacterial heme stores and minimise external stressors that may influence the outcome of competitive studies (29–31). Bacterial suspensions of ~1.0 OD_600_ were made in TSB from 8–10 hr growth on CA and diluted 1:10 in 5 pre-warmed sTSB (5 mL). Broths were incubated for 12 hr at 37°C aerobically with shaking (220 RPM), centrifuged at 4000 × g for 5 min at 37°C and resuspended in fresh, pre-warmed TSB to an OD_600_ of 1.0 prior to use in growth experiments.

### Determination of NTHi-inhibitory activity

A well diffusion assay of the extracted HPL protein was used to categorise Hh strains containing the *HPL* ORF as either HPL-producers (Hh-HPL^+^) or non-producers (Hh-HPL^−^), as previously described (28). This assay was also used to establish the relative inhibitory activity of each strain. Testing was conducted on two indicator NTHi strains (ATCC 49427 and clinical isolate NTHi-L15). Two Hh strains were included as additional *HPL* ORF negative controls for HPL extraction: Hh ATCC 33390 and BW1^*HPL−*^.

### Triplex real-time PCR for the quantification of NTHi, Hh and detection of *HPL*

A real-time quantitative triplex PCR assay was designed to quantify NTHi, Hh and detect the *HPL* open reading frame (ORF). The targets used for discrimination of Hh (*hypD*) and NTHi (*siaT*) have previously been described and validated (32). For detection of the *HPL* ORF, primers were designed based on the *HPL* ORF of Hh-BW1 (29) (GenBank MN720274). The FAM, HEX and TET channels were used for simultaneous fluorescence detection of *siaT*, *hypD* and *HPL*, respectively. Primer and probe sequences are detailed in Table 1. Primer specificity was confirmed by discontiguous megaBLAST analysis and PCR of a panel of *Haemophilus spp.* and multiple genera representing common upper respiratory tract flora. PCR assays were extensively optimised and evaluated for detection/quantification limits in triplex format.

**Table 1.**
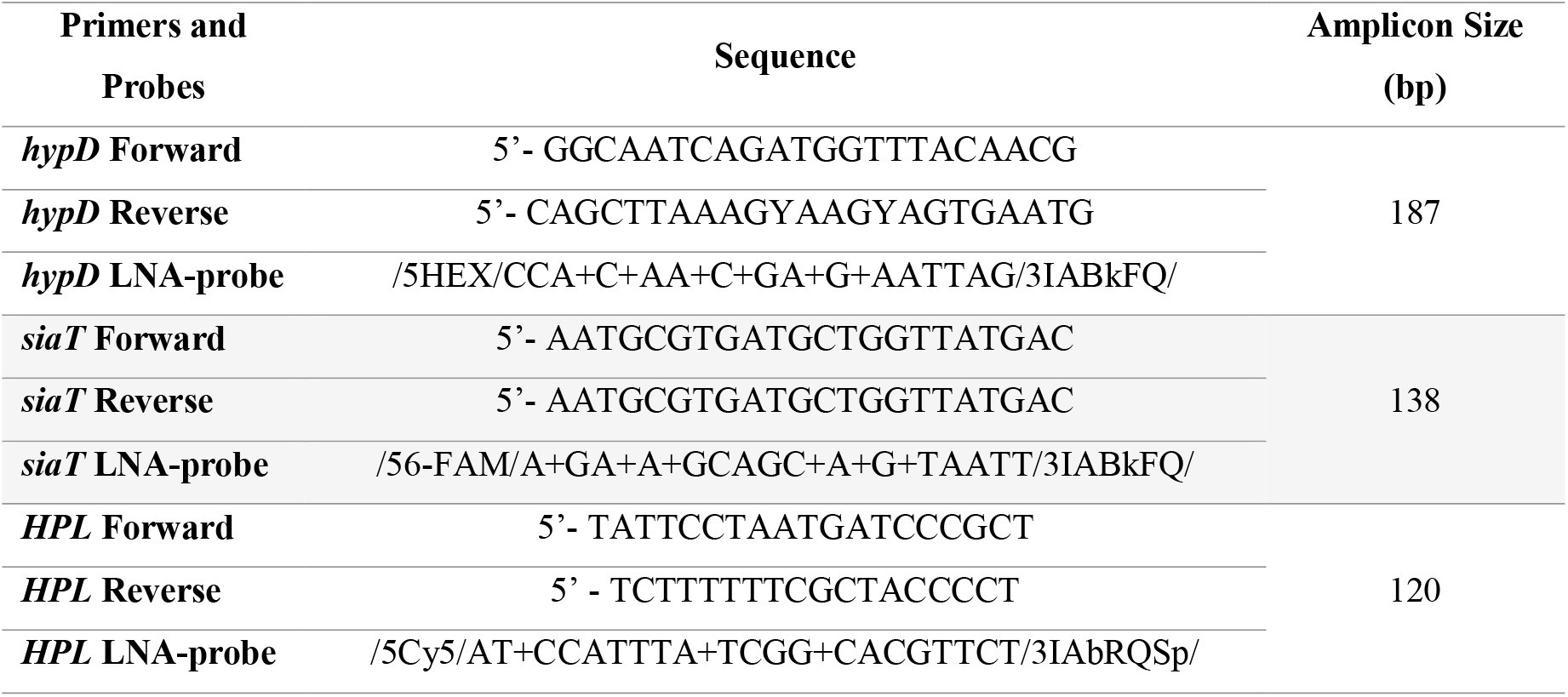
Summary of primer and LNA-probe sequences, and expected amplicon size for the *hypD*, *siaT and HPL* targets.

PCRs were performed using the CFX96 Touch™ real-time PCR system (Bio-Rad) in 96-well optical plates. Polymerase activation was performed at 95°C for 3 minutes, followed by 40 amplification cycles of denaturation at 95°C for 15 seconds, and annealing at 62°C for 1 minute. Each reaction contained 0.25 μM of *hypD, siaT* and *HPL* primers, 0.1 μM LNA-probes, 1× PrimeTime master mix (Integrated DNA Technologies) and 5 μl genomic DNA and molecular-grade water, to a total volume of 20 μl. Template DNA was prepared by a thermal extraction protocol and tested in duplicate. Each run included a positive control for the *HPL* ORF (Hh-BW1), negative control (*H. parainfluenzae* ATCC 7901), no-template control and 10-fold dilutions of a standard containing 2 × 10^−8^ ng NTHi ATCC 49247 and Hh ATCC 33390 DNA. Quantification of NTHi and Hh was expressed as genome equivalents (GE) calculated from the standard, as previously described (32). Bacterial quantification from thermally extracted DNA was validated against conventional quantification by optical density and colony counts. Complete details of PCR primer design, assay optimisation and DNA extraction protocol evaluation are available in supplementary material.

### Competition Assays

#### Short-term broth competition

Culture mixes were prepared by adding 100 μL of heme-replete preparations of Hh-HPL^+^ or Hh-HPL^−^ and 100 μL of NTHi ATCC 49247 to 5 mL pre-warmed sTSB containing 0.0, 0.9, 3.8 or 15.0 ug mL^−1^ porcine hematin. Broths containing single strains were also prepared in parallel to determine baseline growth. Broths were incubated aerobically on an incubator shaker at 37°C (220 RPM) for 16 hours. At different time intervals, aliquots of 50 μL were taken for boiled gDNA extraction and subsequent triplex PCR quantification of GE. Aliquots of 500 μL were taken at 8 hours for quantification of *HPL* expression. Purity of broth growth was checked by plating on CA after 16 hours incubation.

Statistical comparisons were made between strains grown with a competitor and baseline growth by calculating the change in the number of cells per hour (growth rate) using the following formula:

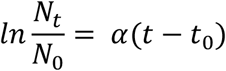

Where ***N*_*t*_** is the number of cells (measured as GE) at time ***t***, *N*_0_ is the number of cells at time zero (***t***_0_), and ***α*** is the growth rate where units are determined by the units of ***t***.

#### Fitness assay

Culture mixes were prepared by adding 100 μL of heme-replete preparations of Hh-HPL^+^ or Hh-HPL^−^ and 100 μL of NTHi (ATCC 49247, ATCC 49766 or NCTC 11315) to 5 mL of pre-warmed sTSB containing 0.0, 0.9, 3.8 or 15.0 μg mL^−1^ porcine hematin. Broths were incubated aerobically on an incubator shaker at 37°C (220 RPM) for 12 hours prior to sub-culture (200 μL) in fresh sTSB (2 mL) containing the same concentration of heme as the inoculum. The process of 12-hourly incubation followed by sub-culture into fresh broth was repeated until 6 days had elapsed. After each 12-hour incubation, aliquots of 50 μL were taken for boiled gDNA extraction and subsequent triplex PCR quantification of GE. Purity of broth growth was confirmed by plating on CA after each 12-hour incubation. Fitness of NTHi at each time point was determined using the following equation (33):

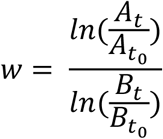

Where ***w*** is fitness, **A** and **B** are the population sizes of the two competitors, subscripts ***t*_*0*_** and ***t*** indicate the initial and final time points in the assay. Growth after the first 12-hour culture was used as baseline for fitness determination (***t*_*0*_**).

### Expression analysis

#### RNA extraction, purification and quantification

Aliquots taken from broth growth were immediately added to two volumes of RNAprotect Bacteria Reagent (Qiagen) for immediate stabilization of bacterial RNA. Stabilized aliquots were normalized to an OD_600_ of 0.05 (approximately 5 × 10^7^ cells), pelleted by centrifugation for 10 minutes at 5000 × g and stored at −20°C overnight. Bacterial lysates were prepared by resuspending pellets in 100 μl TE buffer (30mM Tris-Cl, 1 mM EDTA, pH 8.0) containing 15 mg mL^−1^ lysozyme and 20 μL proteinase K, vortexed and incubated at room temperature in an incubator shaker (1000 RPM) for 1 hr. Following addition of 350 μL RLT buffer, samples were vortexed and centrifuged at 20000 × g for 2 minutes. The supernatant was purified following the manufacturers protocol for RNeasy Plus Mini Kit which was semi-automated by the QIAcube (Qiagen). The iScript™ cDNA Synthesis Kit (Bio-Rad) was used to produce cDNA for subsequent PCR. The validated triplex PCR was used to determine expression of *HPL* ORF in Hh strains, using *hypD* as the housekeeper gene.

#### Expression validation

Expression analysis was employed to determine baseline expression and suitability of prospective competitive test conditions for *HPL* expression. Given the kinetics of bacterial growth and the heme-binding capacity of HPL, time and heme availability were targeted as factors that may influence *HPL* expression. The *hypD* target was selected as a potential housekeeper gene and validated for test conditions.

Heme-starved preparations of Hh-BW1 and the Hh-BW1^*HPL*−^ (100 μL) were added to 5 mL pre-warmed sTSB containing either 0.0 or 15.0 μg ml^−1^ of porcine haematin. Broths were incubated for 8 hours and aliquots of 500 μL were removed for RNA extraction and purification at 0, 4 and 8 hours.

### Statistical analysis

Statistical analysis was performed using GraphPad Prism V7.04, 2017. Statistical significance was determined by comparison of growth data (growth rate or fitness) between strains grown with a competitor and baseline growth. Data were tested for normality using the Shapiro-Wilk test, followed by a two-way ANOVA with Dunnett’s multiple comparison test. Expression ratios and statistical significance were calculated with 2000 iterations by the Relative Expression Software Tool (REST; v 1.0, 2009) (34, 35).

## RESULTS AND DISCUSSION

### Validation of a triplex real-time PCR for quantification of NTHi, Hh and detection of HPL

The *HPL* amplicon was confirmed to be specific and sensitive for the detection of the five previously identified *HPL* sequence variants (29) by *in silico* investigations and by PCR. Specificity of the *HypD* and *SiatT* targets was also confirmed by PCR. Complete results of PCR assay validation is detailed in supplementary materials. The low limit of quantification values for the *HypD* and *SiaT* assays in triplex format were 2 × 10^−5^ ng and 2 × 10^−4^ ng, corresponding to 10 and 100 GE respectively. The lower limit of detection for the *HPL* assay was 10 GE (Figure S1). The upper limits of detection/quantification were not explicitly determined as expected DNA levels from sample were unlikely to exceed the maximum 2 ng tested.

Given the high volume of samples generated from growth experiments, a cheap and high-throughput DNA extraction method was required to reliably distinguish and quantify NTHi and Hh in co-culture. Extraction utilizing thermal lysis has previously been shown to be an efficient and cost-effective method to harvest bacterial DNA for quantitative real-time PCR from suspensions of several bacterial species in a range of sample matrices (36–41). Crude DNA extraction methods are also prone to contamination with PCR inhibitors originating from sample matrices (39, 42). There are also reports of intra- and inter-species differences in DNA extractions efficiencies (39, 42, 43). PCR quantification of gDNA extracted by thermal lysis was validated and found to be comparable to quantification by OD_600_ and colony counts. Full details of thermal extraction validation are available in supplementary materials.

### Baseline NTHi-inhibitory activity of Hh-HPL^+^ strains

We previously discovered a number of distinct genetic variations of the *HPL* ORF with varying NTHi-inhibitory activity, as determined by functional screening (29). The two Hh-HPL^+^ clinical isolates (Hh-BW1 and Hh-RHH122) that exhibited the highest inhibitory capacity share 100% sequence similarity in the *HPL* ORF. Thus, strains containing this sequence variant were selected for further investigation.

A well diffusion assay was employed to confirm inhibitory capacity and establish relative baseline NTHI-inhibitory activity, normalised for population density. Hh-RHH122 and Hh-BW1 demonstrated the highest inhibitory activity, while Hh-NF5 and Hh-NF4 served as intermediate (~50%) and non-inhibitory respectively (Figure S5).

### HPL-production mediates NTHi growth inhibition and a competitive advantage in Hh

The growth rate of NTHi was significantly impaired during competition with all Hh-HPL^+^ strains, compared to growth without competition (p < 0.0001) (Figure 1A). This inhibitory effect was more pronounced during competition with highly bioactive strains (Hh-BW1 and Hh-RHH122), compared to the intermediate (Hh-NF5). The growth rate of NTHi during competition with Hh-HPL^−^ was not significantly affected (Figure 1B, suggesting that the inhibitory effect observed was a characteristic of Hh-HPL^+^ strains.

**Figure 1.**
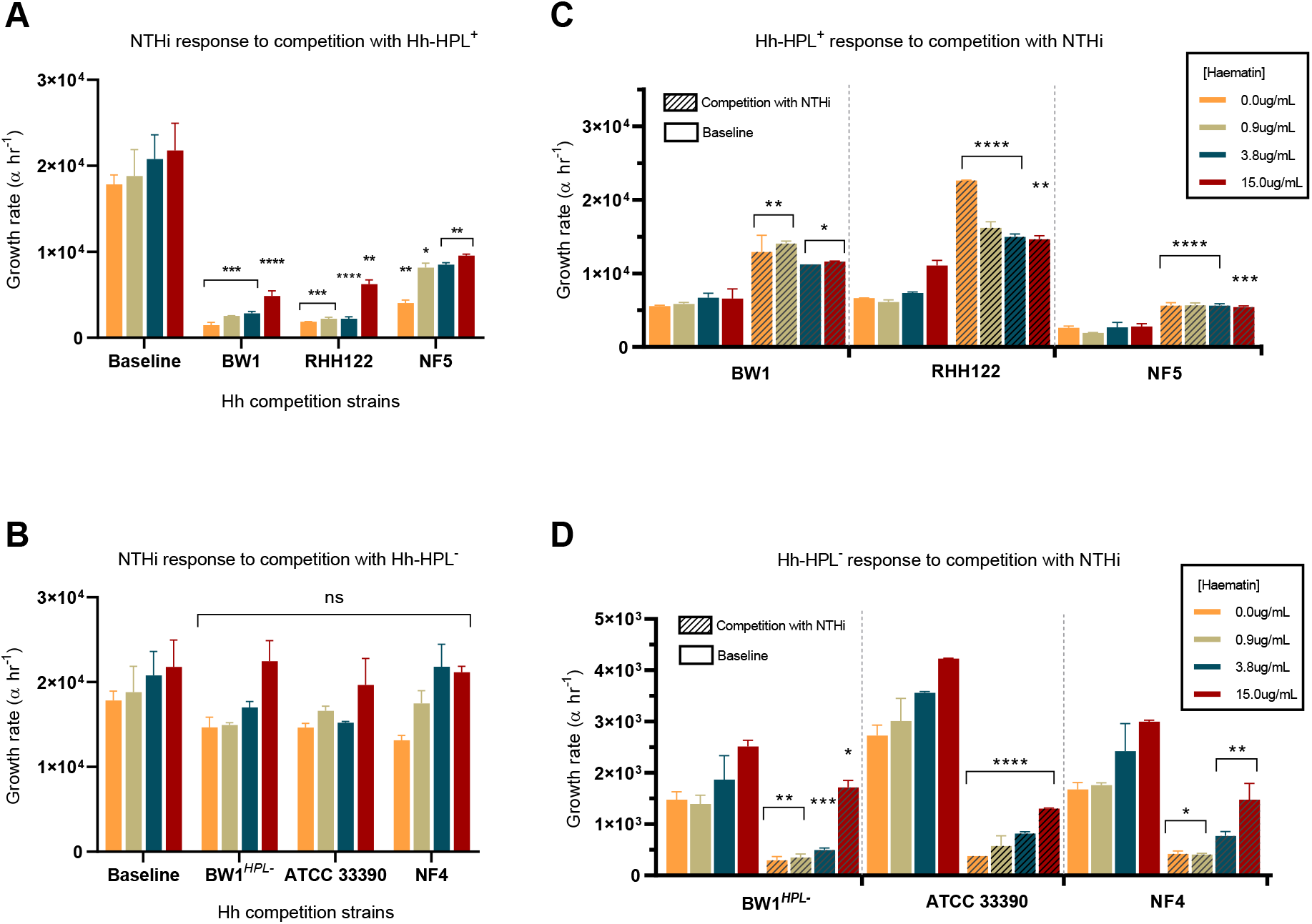
Short-term competition between NTHi and Hh. Calculated growth rates of NTHi in response to competition with **(A)** Hh-HPL^+^ or **(C)** Hh-HPL^−^. The growth rate for each **(B)** Hh-HPL^+^ and **(D)** Hh-HPL^−^ strain was also determined. Data points represented as mean +/− SEM of three separate experiments, performed in triplicate; P < 0.05*, p < 0.005 **, p < 0.0005 ***, p < 0.0001 ****.

For commensals and pathogens living in or invading human tissues, iron-containing heme is often a limiting nutrient, particularly in the respiratory tract where concentrations are considered to be low (44). This is particularly true for heme auxotrophs including NTHi and Hh; for these species survival in the URT niche is dependent on their ability to outcompete host proteins and co-existing bacterial populations for heme (20). We previously demonstrated that the NTHi-inhibitory mechanism of HPL is associated with it’s ability to bind heme in a form inaccessible to NTHi and that inhibitory activity is lost in conditions where heme concentration exceeds the binding capacity of HPL (29). While levels of heme/iron are considered to be low in the respiratory tract, there is indirect evidence for increased heme/iron levels in airways of smokers, COPD and CF which may contribute to increased susceptibility to infection in these individuals (44). Thus, it was important to assess the effectiveness of HPL with varying concentrations of heme to ensure probiotic effectiveness in a range of *in vitro* conditions reflecting possible *in vivo* scenarios. The NTHi-inhibitory capacity of HPL was maintained even in conditions of high heme-availability (15 μg mL^−1^), albeit to a lesser degree than lower heme concentrations (0.0-3.8 μg/mL) (Figure 1A). This suggests that levels of HPL produced by Hh are sufficient to limit NTHi’s access to heme in a dynamic *in vitro* system, even under excess heme conditions unlikely to be encountered *in vivo* (44).

Interestingly, Hh-HPL^+^ strains exhibited a pattern of enhanced growth in response to NTHi competition (p <0.0001) (Figure 1B). This effect was observed in all heme concentrations and was more pronounced in the highly inhibitory Hh-HPL^+^ strains. The converse was observed in Hh-HPL^−^strains, where they exhibited poorer growth in response to competition with NTHi (Figure 1D). This may be a reflection of the highly efficient set of heme-scavenging systems possessed by NTHi that outcompete Hh in the absence of HPL.

### NTHi-inhibitory capacity is associated with expression of HPL

To further test whether the observed competitive advantage of Hh-HPL^+^ strains could be attributed to HPL production, *HPL* expression was quantified by reverse transcription and real-time PCR during competitive growth with NTHi. The *hypD* target was validated as the housekeeper gene (Figure S6A) and the optimal growth phase for *HPL* expression analysis was determined (Figure S6B).

Baseline expression of *HPL* was highest in the highly bioactive strains (Hh-BW1 and Hh-RHH122), significantly lower in Hh-NF5 (p <0.0001), and completely absent in Hh-NF4 (Figure 2A). These results establish a connection between expression of HPL and NTHi-inhibitory activity as demonstrated by well-diffusion and short-term competition studies (Figure 1). Upregulation of *HPL* was observed in all Hh-HPL^+^ in response to competition with NTHi, an effect that was more pronounced in Hh-BW1 and Hh-RHH122 (Figure 2B). This may explain the enhanced growth of Hh-HPL^+^ strains in response to NTHi during the short-term competition assays (Figure 1C). These results show that expression of *HPL* has a significant impact on the NTHi-inhibitory capacity of Hh-HPL^+^ strains and therapeutic utility in an *in vivo* setting. Therefore, the huge differential expression of *HPL* amongst Hh-HPL^+^ strains must be considered when selecting a probiotic candidate. However, our understanding of HPL regulation is still rudimentary. Further investigation into potential upstream regulatory components or post-translational modification is needed to elucidate the inter-strain differences in HPL production and/or biological activity despite complete ORF sequence homology.

**Figure 2.**
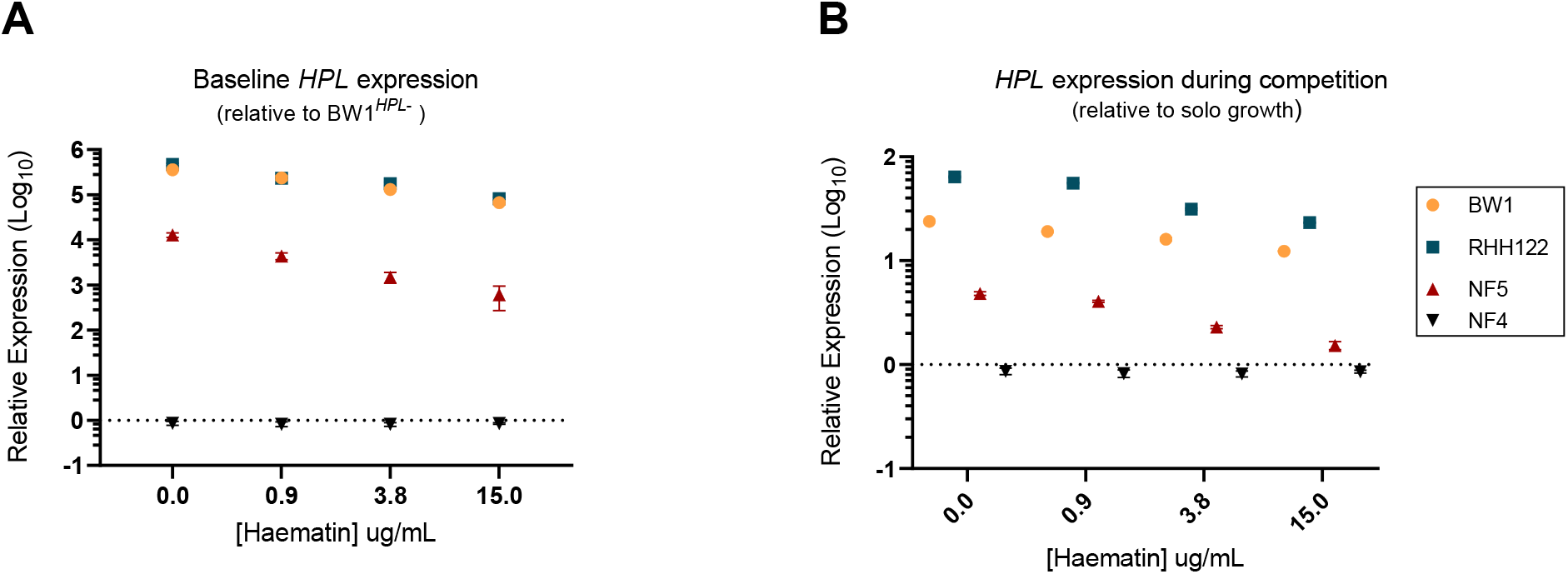
*HPL* expression among Hh strains during competition with NTHi. PCR-quantified expression of *HPL* **(A)** for baseline expression (compared to Hh-BW1^HPL−^) or **(B)** during competition with NTHi relative to individual growth. Data points represented as mean +/− SEM of four biological replicates, performed from duplicate RNA extractions.

### NTHi fitness dramatically decreases in competition with HPL-producers

Short-term competition may highlight the potency of HPL-mediated inhibition but is not representative of *in vivo* competition dynamics. Thus, a longer-term study was employed to assess the competition between NTHi and Hh-HPL^+^ over a period of 6 days (12 generations). The competitive advantage of Hh-HPL+ strains was evident within the 2^nd^ (24 hours) and 4^th^ generations (48 hours) with highly bioactive Hh-HPL^+^, or the intermediate Hh-HPL^+^, respectively (Figure 3A). The stunted inhibitory activity exhibited by Hh-NF5 may be attributed to lower levels of HPL production over the course of the assay. The fitness of NTHi over subsequent generations decreases significantly until complete loss of fitness during the final generations. Although, competition with Hh-HPL^−^ did not result in a significant loss of fitness over the 6-day period, there was decrease in fitness of all Hh-HPL strains at 24 hours, followed by complete recovery (Figure 3A). This may have arisen from competition for heme prior to the onset of maximum HPL production.

**Figure 3.**
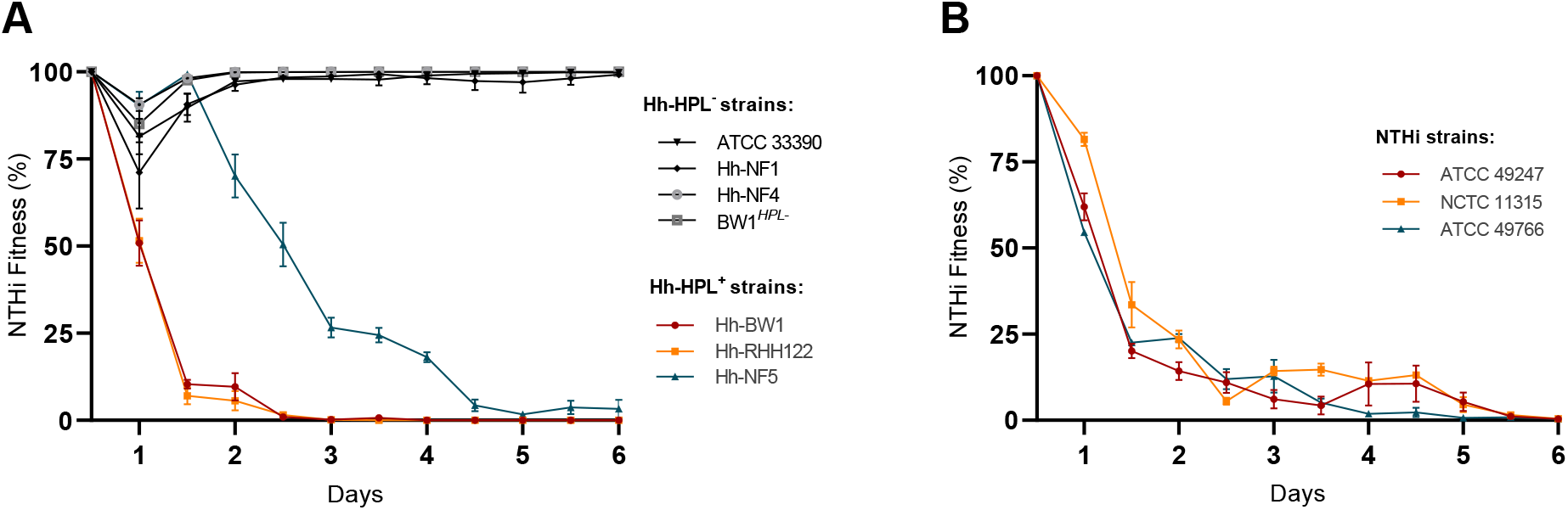
Fitness of NTHi strains during co-culture with Hh. Calculated fitness of NTHi in response to competition with Hh-HPL^+^ or Hh-HPL^−^ relative to growth of the competitor strain. **(A)** Competition betweeb a single NTHi strain and multiple Hh, or **(B)** multiple NTHi against Hh-BW1. Data points represented as mean +/− SEM of three separate experiments, performed in quadruplicate.

To show that loss of fitness of NTHi was not unique to NTHi strain ATCC 49247, additional reference strains NCTC 11315 and ATCC 49766 were tested in competition with Hh-BW1. All three NTHi strains responded in the same manner, culminating in a total loss of fitness at the end of the 6-day period (Figure 3B).

## CONCLUSION

Previously, we identified an uncharacterized hemophore, designated hemophilin, produced by Hh which is able to inhibit NTHi growth by heme starvation (29). The current study aimed to further test the inhibitory capacity of Hh-HPL^+^ by direct *in vitro* competition with NTHi, for the purpose of determining probiotic potential.

The unique inhibitory capacity and growth advantage exhibited by Hh-HPL^+^ strains, together with expression analysis during competitive growth confirmed that inhibition was mediated by the presence of HPL. These results demonstrate the enormous probiotic potential Hh-HPL^+^ strains against NTHi colonization in the URT. Reduction or elimination of NTHi carriage from the URT and subsequent migration to the lower airways would be an effective means of preventing infection with the organism. Further investigation is required to determine the protective capacity of HPL-producers *in vivo* and their ability to interrupt NTHi colonization of host cells.

## REFERENCES

1. Mukundan D, Ecevit Z, Patel M, Marrs CF, Gilsdorf JR. 2007. Pharyngeal colonization dynamics of Haemophilus influenzae and Haemophilus haemolyticus in healthy adult carriers. Journal of clinical microbiology 45:3207–3217.

2. Puig C, Grau I, Marti S, Tubau F, Calatayud L, Pallares R, Liñares J, Ardanuy C. 2014. Clinical and molecular epidemiology of Haemophilus influenzae causing invasive disease in adult patients. PLoS One 9:e112711.

3. Murphy TF, Faden H, Bakaletz LO, Kyd JM, Forsgren A, Campos J, Virji M, Pelton SI. 2009. Nontypeable Haemophilus influenzae as a pathogen in children. The Pediatric infectious disease journal 28:43–48.

4. Slack MP. 2015. A review of the role of Haemophilus influenzae in community-acquired pneumonia. Pneumonia 6:26–43.

5. Murphy TF. 2015. Vaccines for nontypeable Haemophilus influenzae: the future is now. Clin Vaccine Immunol 22:459–466.

6. King P. 2012. Haemophilus influenzae and the lung (Haemophilus and the lung). Clinical and translational medicine 1:1–9.

7. Langereis JD, de Jonge MI. 2015. Invasive disease caused by nontypeable Haemophilus influenzae. Emerging infectious diseases 21:1711.

8. van Wessel K, Rodenburg GD, Veenhoven RH, Spanjaard L, van der Ende A, Sanders EA. 2011. Nontypeable Haemophilus influenzae invasive disease in The Netherlands: a retrospective surveillance study 2001–2008. Clinical Infectious Diseases 53:e1–e7.

9. Giufrè M, Fabiani M, Cardines R, Riccardo F, Caporali MG, D’Ancona F, Pezzotti P, Cerquetti M. 2018. Increasing trend in invasive non-typeable Haemophilus influenzae disease and molecular characterization of the isolates, Italy, 2012–2016. Vaccine 36:6615–6622.

10. Sriram KB, Cox AJ, Clancy RL, Slack MP, Cripps AW. 2017. Nontypeable Haemophilus influenzae and chronic obstructive pulmonary disease: a review for clinicians. Critical Reviews in Microbiology:1–18.

11. Maddi S, Kolsum U, Jackson S, Barraclough R, Maschera B, Simpson KD, Pascal TG, Durviaux S, Hessel EM, Singh D. 2017. ampicillin resistance in Haemophilus influenzae from COPD patients in the UK. International journal of chronic obstructive pulmonary disease 12:1507.

12. Desai H, Richter S, Doern G, Heilmann K, Dohrn C, Johnson A, Brauer A, Murphy T, Sethi S. 2010. Antibiotic resistance in sputum isolates of Streptococcus pneumoniae in chronic obstructive pulmonary disease is related to antibiotic exposure. COPD: Journal of Chronic Obstructive Pulmonary Disease 7:337–344.

13. Pettigrew MM, Tsuji BT, Gent JF, Kong Y, Holden PN, Sethi S, Murphy TF. 2016. Effect of fluoroquinolones and macrolides on eradication and resistance of Haemophilus influenzae in chronic obstructive pulmonary disease. Antimicrobial agents and chemotherapy 60:4151–4158.

14. Wilson R, Sethi S, Anzueto A, Miravitlles M. 2013. Antibiotics for treatment and prevention of exacerbations of chronic obstructive pulmonary disease. Journal of Infection 67:497–515.

15. Puig C, Tirado-Vélez JM, Calatayud L, Tubau F, Garmendia J, Ardanuy C, Marti S, Adela G, Liñares J. 2015. Molecular characterization of fluoroquinolone resistance in nontypeable Haemophilus influenzae clinical isolates. Antimicrobial agents and chemotherapy 59:461–466.

16. Vila J, Ruiz J, Sanchez F, Navarro F, Mirelis B, de Anta MTJ, Prats G. 1999. Increase in Quinolone Resistance in aHaemophilus influenzae Strain Isolated from a Patient with Recurrent Respiratory Infections Treated with Ofloxacin. Antimicrobial agents and chemotherapy 43:161–162.

17. Bastida T. 2003. Levofloxacin treatment failure in Haemophilus influenzae pneumonia.

18. Hariadi NI, Zhang L, Patel M, Sandstedt SA, Davis GS, Marrs CF, Gilsdorf JR. 2015. Comparative profile of heme acquisition genes in disease-causing and colonizing nontypeable Haemophilus influenzae and Haemophilus haemolyticus. Journal of clinical microbiology 53:2132–2137.

19. Szelestey BR, Heimlich DR, Raffel FK, Justice SS, Mason KM. 2013. Haemophilus responses to nutritional immunity: epigenetic and morphological contribution to biofilm architecture, invasion, persistence and disease severity. PLoS pathogens 9:e1003709.

20. Morton DJ, Bakaletz LO, Jurcisek JA, VanWagoner TM, Seale TW, Whitby PW, Stull TL. 2004. Reduced severity of middle ear infection caused by nontypeable Haemophilus influenzae lacking the hemoglobin/hemoglobin–haptoglobin binding proteins (Hgp) in a chinchilla model of otitis media. Microbial pathogenesis 36:25–33.

21. Morton DJ, Seale TW, Bakaletz LO, Jurcisek JA, Smith A, VanWagoner TM, Whitby PW, Stull TL. 2009. The heme-binding protein (HbpA) of Haemophilus influenzae as a virulence determinant. International Journal of Medical Microbiology 299:479–488.

22. Seale TW, Morton DJ, Whitby PW, Wolf R, Kosanke SD, VanWagoner TM, Stull TL. 2006. Complex role of hemoglobin and hemoglobin-haptoglobin binding proteins in Haemophilus influenzae virulence in the infant rat model of invasive infection. Infection and immunity 74:6213–6225.

23. Ahearn CP, Gallo MC, Murphy TF. 2017. Insights on persistent airway infection by non-typeable Haemophilus influenzae in chronic obstructive pulmonary disease. Pathogens and Disease 75.

24. Stites SW, Plautz MW, Bailey K, O’Brien-Ladner AR, Wesselius LJ. 1999. Increased concentrations of iron and isoferritins in the lower respiratory tract of patients with stable cystic fibrosis. American journal of respiratory and critical care medicine 160:796–801.

25. White DC, Granick S. 1963. Hemin biosynthesis in Haemophilus. Journal of bacteriology 85:842–850.

26. Sgheiza V, Novick B, Stanton S, Pierce J, Kalmeta B, Holmquist MF, Grimaldi K, Bren KL, Michel LV. 2017. Covalent bonding of heme to protein prevents heme capture by nontypeable Haemophilus influenzae. FEBS open bio 7:1778–1783.

27. Skaar EP. 2010. The battle for iron between bacterial pathogens and their vertebrate hosts. PLoS pathogens 6:e1000949.

28. Latham RD, Gell DA, Fairbairn RL, Lyons AB, Shukla SD, Cho KY, Jones DA, Harkness NM, Tristram SG. 2017. An isolate of Haemophilus haemolyticus produces a bacteriocin-like substance that inhibits the growth of nontypeable Haemophilus influenzae. International journal of antimicrobial agents 49:503–506.

29. Latham RD, Torrado M, Atto B, Walshe JL, Wilson R, Guss JM, Mackay JP, Tristram S, Gell DA. 2019. A heme-binding protein produced by Haemophilus haemolyticus inhibits non-typeable Haemophilus influenzae. Molecular Microbiology, In press doi:10.1101/626416:626416.

30. Vogel AR, Szelestey BR, Raffel FK, Sharpe SW, Gearinger RL, Justice SS, Mason KM. 2012. SapF-mediated heme-iron utilization enhances persistence and coordinates biofilm architecture of *Haemophilus*. Front Cell Infect Microbiol 2:42.

31. Mason KM, Raffel FK, Ray WC, Bakaletz LO. 2011. Heme utilization by nontypeable *Haemophilus influenzae* is essential and dependent on Sap transporter function. J Bacteriol 193:2527–35.

32. Price EP, Harris TM, Spargo J, Nosworthy E, Beissbarth J, Chang AB, Smith-Vaughan HC, Sarovich DS. 2017. Simultaneous identification of Haemophilus influenzae and Haemophilus haemolyticus using real-time PCR. Future Microbiology.

33. Wiser MJ, Lenski RE. 2015. A comparison of methods to measure fitness in Escherichia coli. PLoS One 10:e0126210.

34. Pfaffl MW. 2001. A new mathematical model for relative quantification in real-time RT–PCR. Nucleic acids research 29:e45–e45.

35. Pfaffl MW, Horgan GW, Dempfle L. 2002. Relative expression software tool (REST©) for group-wise comparison and statistical analysis of relative expression results in real-time PCR. Nucleic acids research 30:e36–e36.

36. Reischl U, Linde H-J, Metz M, Leppmeier B, Lehn N. 2000. Rapid identification of methicillin-resistantStaphylococcus aureus and simultaneous species confirmation using real-time fluorescence PCR. Journal of Clinical Microbiology 38:2429–2433.

37. Reischl U, Pulz M, Ehret W, Wolf HJ. 1994. PCR-based detection of mycobacteria in sputum samples using a simple and reliable DNA extraction protocol. BioTechniques 17:844–845.

38. Sweeney RW, Whitlock RH, McAdams SC. 2006. Comparison of three DNA preparation methods for real-time polymerase chain reaction confirmation of Mycobacterium avium subsp. paratuberculosis growth in an automated broth culture system. Journal of veterinary diagnostic investigation 18:587–590.

39. Van Tongeren S, Degener J, Harmsen H. 2011. Comparison of three rapid and easy bacterial DNA extraction methods for use with quantitative real-time PCR. European journal of clinical microbiology & infectious diseases 30:1053–1061.

40. Wilson DA, Yen-Lieberman B, Reischl U, Gordon SM, Procop GW. 2003. Detection of Legionella pneumophila by real-time PCR for the mip gene. Journal of clinical microbiology 41:3327–3330.

41. Freschi CR, Oliveira CJBd. 2005. Comparison of DNA-extraction methods and selective enrichment broths on the detection of Salmonella Typhimurium in swine feces by polymerase chain reaction (PCR). Brazilian Journal of Microbiology 36:363–367.

42. Coyne SR, Craw PD, Norwood DA, Ulrich MP. 2004. Comparative analysis of the Schleicher and Schuell IsoCode Stix DNA isolation device and the Qiagen QIAamp DNA mini kit. Journal of clinical microbiology 42:4859–4862.

43. Rantakokko-Jalava K, Jalava J. 2002. Optimal DNA isolation method for detection of bacteria in clinical specimens by broad-range PCR. Journal of clinical microbiology 40:4211–4217.

44. Ali MK, Kim RY, Karim R, Mayall JR, Martin KL, Shahandeh A, Abbasian F, Starkey MR, Loustaud-Ratti V, Johnstone D. 2017. Role of iron in the pathogenesis of respiratory disease. The international journal of biochemistry & cell biology 88:181–195.

